# Synergistic Activity of Silver Nanoparticles and Vancomycin Against a Spectrum of *Staphylococcus aureus* Biofilm Types

**DOI:** 10.1101/337436

**Authors:** Bryan B. Hair, Matthew E. Conley, Trevor M. Wienclaw, Mark J. Conley, Bradford K. Berges

## Abstract

*Staphylococcus aureus* (SA) is an important human pathogen, causing potentially lethal infections of the blood, skin, and lungs. SA is becoming increasingly difficult to treat due to high levels of antibiotic resistance, and new treatments are needed. SA is able to evade antibiotics and immune surveillance through biofilm development. Biofilms are communities of microorganisms that are able to prevent the entry of antimicrobials and immune cells. Biofilms are also involved in SA transmission because biofilms can form on medical devices. In this study, we tested silver nanoparticles and vancomycin for their anti-biofilm effects on SA. We used 10 different SA isolates, representing a spectrum of biofilm-forming ability, and a crystal violet assay to measure biofilm mass. 2μg/mL vancomycin treatment resulted in a significant reduction in established SA biofilms in 7/10 isolates, including 4/5 methicillin susceptible SA (MSSA) and 3/5 methicillin-resistant (MRSA) isolates (mean reduction in crystal violet stain of 13.0%; high of 26.5% and low of 0%). Silver nanoparticle treatment of SA biofilms resulted in a significant reduction in 6/10 isolates, including 4/5 MSSA and 2/5 MRSA (mean reduction of 8.7%; high of 21.2% and low of 0%). A combinatorial treatment with silver nanoparticles and vancomycin resulted in significant reductions in 9/10 isolates (mean reduction of 20.8%; high of 39.3% and low of 0%). We conclude that both vancomycin and silver nanoparticle treatment of established tissue culture-based SA biofilms result in significant reductions in biofilm mass, with a combinatorial treatment even more effective than either treatment alone.

## Introduction

*Staphylococcus aureus* (SA) is a widespread commensal bacterium that, because of its pathogenicity and ability to persist in hospital settings, has become a global healthcare concern. This gram positive coccus colonizes the anterior nares of roughly 30% of the population, and several of its characteristics allow it to become virulent and disseminate effectively through the host (1). Epidemiological reports estimate that there are annually 477,927 hospitalizations and 29,164 deaths that were related to SA in the United States alone (2). In addition to antibiotic resistance, the ability to form a community of cells, or biofilm, is a virulence factor that can promote transmission and exacerbate disease.

Bacterial biofilms are ubiquitous and likely the predominant form in which most prokaryotes exist in nature (3). As bacterial cells grow collectively, they are able to create a protective extracellular matrix and initiate other processes that ultimately increase their viability. Potential benefits include amplified resistance to bactericidal agents such as antibiotics, bacteriophage, and host immune defenses (4). Often, multiple bacterial species are incorporated into a single biofilm, and this can facilitate horizontal gene transfer (5). *S. aureus* has numerous proteins on its outer membrane (Microbial Surface Components Recognizing Adhesive Matrix Molecules) that allow cells to adhere to a wide variety of surfaces (6). Once adhered, a distinct set of genes become activated, resulting in the formation of a biofilm (7). Exopolysaccharides, proteins, extracellular DNA (eDNA), and other polymers have been identified as some of the important constituents of the extracellular matrix that promote intercellular adhesion (8, 9). Although there is a spectrum of biofilm types in SA, in general they are often characterized as being either polysaccharide-based or eDNA and protein-based (10).

The capacity to form robust biofilms is clinically relevant. Although there are countless SA isolates that have arisen from the community and entered hospitals, there are currently only five major clonal lineages of *S. aureus* that account for the majority of nosocomial infections at a global level (11). When one such strain arose in Brazil during the 1990s, it was discovered that it had a much greater ability to produce biofilms and to adhere to polystyrene surfaces (in addition to being multi-drug resistant, a property shared by other strains) (6, 12). It could thereby effectively colonize prosthetic devices, catheters, and damaged tissues.

Antibiotics have saved millions of lives, but their common use has resulted in widespread antibiotic resistance. Within one year of the introduction of methicillin as an antibiotic, resistant strains of SA had already been discovered (methicillin-resistant SA or MRSA) (13). SA is equipped with a mobile genetic element known as the staphylococcal cassette chromosome (SCC) mec. SCCmec can integrate into the bacterial chromosome and promotes the capture and sharing of foreign DNA segments between SA isolates and even other species of bacteria (6) (14). SCCmec often carries a gene that encodes β-lactam resistance (*mecA*), in addition to numerous other resistance genes against non-β-lactam antibiotics (15). In the short time since the first reported MRSA, these genes have proliferated and become increasingly common. Based on isolates gathered from intensive care units, roughly 65% of the strains currently encountered in hospitals have a copy of *mecA* and are resistant to methicillin (16).

Vancomycin is now one of our final recourses for treating infections caused by multi-drug resistant SA. In 1996, strains of MRSA from Japan were the first recorded isolates that exhibited resistance to vancomycin (15, 17). Another independently evolved strain emerged in 2002 that resulted from the transfer of the *vanA* gene from enterococci, which conferred vancomycin resistance to SA (18). These strains pose an unsettling dilemma to physicians as there are now no widely accepted treatment options available for these strains. Furthermore, vancomycin is a large glycopeptide and because of its size is less effective at penetrating exopolysaccharides, like those created by the intercellular adhesion gene cluster in staphylococci (19). Similarly, the bactericidal mechanism only acts upon dividing cells, and because cell growth is decreased in biofilms, its potency and effect are marginalized (20, 21). For these reasons, even susceptible SA isolates can be difficult to treat and are requiring greater concentrations of antibiotic to achieve the same effect (22). In comparison to β-lactam antibiotics, vancomycin is associated with higher rates of treatment failure, persistent bacteremia, clinical relapse, and toxicity to the host (23), making it a less than ideal last resort.

The drawbacks of vancomycin, and antibiotics in general, have spurred investigations for possible alternatives. Silver nanoparticles are receiving attention and may show promise as an antibiotic because of their ability to inhibit and eliminate the growth of a wide array of bacteria, including SA (24). The nanoparticles release Ag^+^ ions into solution, which in turn interact with water to create reactive oxygen species (ROS). ROS are able to penetrate biofilms and cell membranes, and subsequent to their entry, they damage cellular components resulting in cell death (25). Human cells have a greater ability to deal with ROS, so at low concentrations silver nanoparticles may be safe for use in humans (26). The mechanisms utilized by silver nanoparticles to kill bacteria are highly conserved and operate on multiple systems; this suggests that the acquisition of resistance is unlikely (27). It should be noted, however, that such agents require high concentrations to achieve their desired effects, and they are able to eliminate bacterial growth to a limited extent (24, 28, 29).

Innovative researchers have sought to use silver nanoparticles in conjunction with antibiotics to increase their potency. Though the mechanism behind their interactions has yet to be fully elucidated, there is a clear synergy when they are used in tandem (30, 31). A recent study went one step further to determine if this synergy would also prevail against established biofilms. They successfully demonstrated that biofilms composed of *Pseudomonas aeruginosa* synergistically reduced by aztreonam and silver nanoparticles (32). No current studies have examined if, and to what level, this synergy is present between vancomycin and silver nanoparticles on SA.

The aim of the present study was to examine the susceptibility of SA biofilms to vancomycin and silver nanoparticles. We first characterized each isolate’s biofilm type via a crystal violet stain, either untreated or with proteinase or DNase treatments intended to identify components in the biofilm. We chose a spectrum of SA isolates, then silver nanoparticles and vancomycin were tested individually for anti-biofilm activity against a number of MRSA and MSSA strains, including those from hospital isolates and from strains isolated in the community. Next, silver nanoparticles and vancomycin were tested together to determine if they produced synergistic results against SA biofilms.

## Results

### Strain characterization for biofilm potential

A number of different SA isolates were analyzed for biofilm-forming capacities and susceptibility to biofilm disruption in these studies. Various attributes of these isolates are presented in Table 1, including the source of the strain, susceptibility/resistance to methicillin, presence of the *icaD* gene (involved in polysaccharide biofilm formation), colony morphology on congo red agar (indicates slime/polysaccharide production), baseline biofilm mass (via crystal violet stain), and biofilm susceptibility to proteinase K and DNase treatment. A commonly used crystal violet biofilm assay was employed in these studies (see Methods).

**Table 1:** *Staphylococcus aureus* strains used in this study.

| Strain name | Source | Susceptibility to methicillin | Congo red colony morphology | *icaD* genotype | Base biofilm mass, OD595 | % Biofilm reduction by proteinase K treatment | % Biofilm reduction by DNase treatment |
| --- | --- | --- | --- | --- | --- | --- | --- |
| M1 | ATF | MRSA | +++ | + | 2.75 +/- 0.61 | 43.1 +/- 1.2% | 42.3 +/- 3.8% |
| M6 | ATF | MRSA | + | - | 3.40 +/- 0.12 | 43.7 +/- 0.8% | 51.3 +/- 2.3% |
| M7 | ATF | MRSA | +++ | - | 2.92 +/- 0.21 | 44.1 +/- 0.8% | 58.0 +/- 2.3% |
| HA1 | Hospital isolate | MRSA | +++ | + | 2.37 +/- 0.08 | 61.2 +/- 13.7% | No reduction |
| USA300 GA-92 | BEI Resources | MRSA | ++ | + | 1.28 +/- 0.10 | 67.8 +/- 8.8% | 33.2 +/- 3.7% |
| SA 29213 | ATCC | MSSA | + | + | 1.24 +/- 0.11 | 35.1 +/- 9.2% | 32.9 +/- 7.3% |
| SA 12600 | ATCC | MSSA | +++ | + | 2.26 +/- 0.10 | 68.3 +/- 9.3% | 34.4 +/- 7.3% |
| SA 4651 | ATCC | MSSA | +++ | + | 0.99 +/- 0.06 | 59.5 +/- 5.7% | 48.0 +/- 7.4% |
| SA 25923 | ATCC | MSSA | +++ | + | 1.28 +/- 0.13 | No reduction | 54.6 +/- 6.7% |
| SA 6538 | ATCC | MSSA | ++ | - | 0.51 +/- 0.031 | 36.05 +/- 8.3% | 46.0 +/- 3.6% |

Strains used in this study included both MRSA and MSSA isolates. ATF=athletic training
facility in Utah, BEI Resources (Biodefense and Emerging Infections Research Resources
Repository), ATCC (American Type Culture Collection). Standard error is indicated.

Overall, 5 MRSA strains and 5 MSSA strains were characterized for biofilm development and composition, including 6 isolates from ATCC or BEI Resources that are publicly available. Most isolates were positive for the *icaD* gene (7/10), and colony morphology on congo red agar was also assessed (6/10 isolates scored as+++). The baseline biofilm mass in untreated samples treatment) varied amongst the isolates examined, with an OD595nm range of 0.51+/-0.031 to 3.40+/-0.12. Overall, a breadth of SA isolates was examined in an effort to represent potential SA strains in the environment.

Treatment of biofilms with proteinase K or DNase can reveal the relative contribution of extracellular proteins or DNA to the structure of a biofilm. We found that some biofilms were similarly affected by either proteinase K or DNase treatment (6/10 total isolates, including 3/5 MRSA and 3/5 MSSA). Some biofilms were more affected by proteinase K than by DNase (3/10 total isolates, including 2/5 MRSA and 1/5 MSSA), and that one isolate was more affected by DNase than by proteinase K (a MSSA). One MRSA isolate was completely resistant to DNase treatment (HA1), and one MSSA isolate was completely resistant to proteinase K treatment (SA 25923).

### Evaluation of vancomycin for anti-biofilm activity

Vancomycin is commonly employed to treat SA infections, since resistance to this antibiotic is typically low. In order to determine the anti-biofilm effects of vancomycin, biofilms were established in tissue culture-treated wells for 24h and then 2μg/mL vancomycin was added (or ddH_2_O as an untreated control; see Fig. 1). We found significant reductions in biofilm staining in 7/10 isolates, with a mean reduction in biofilm mass of 13.0+/-2.7% (p = 0.0009). MRSA isolates and MSSA isolates were similar in their susceptibility to vancomycin, with 3/5 MRSA isolates susceptible (mean reduction of 12.3+/-5.1%; p = 0.04) and 4/5 MSSA isolates susceptible (mean reduction of 13.6+/-2.4%; p = 0.006). There was no significant difference in MRSA vs MSSA susceptibility to vancomycin (p = 0.38).

**Figure 1:**
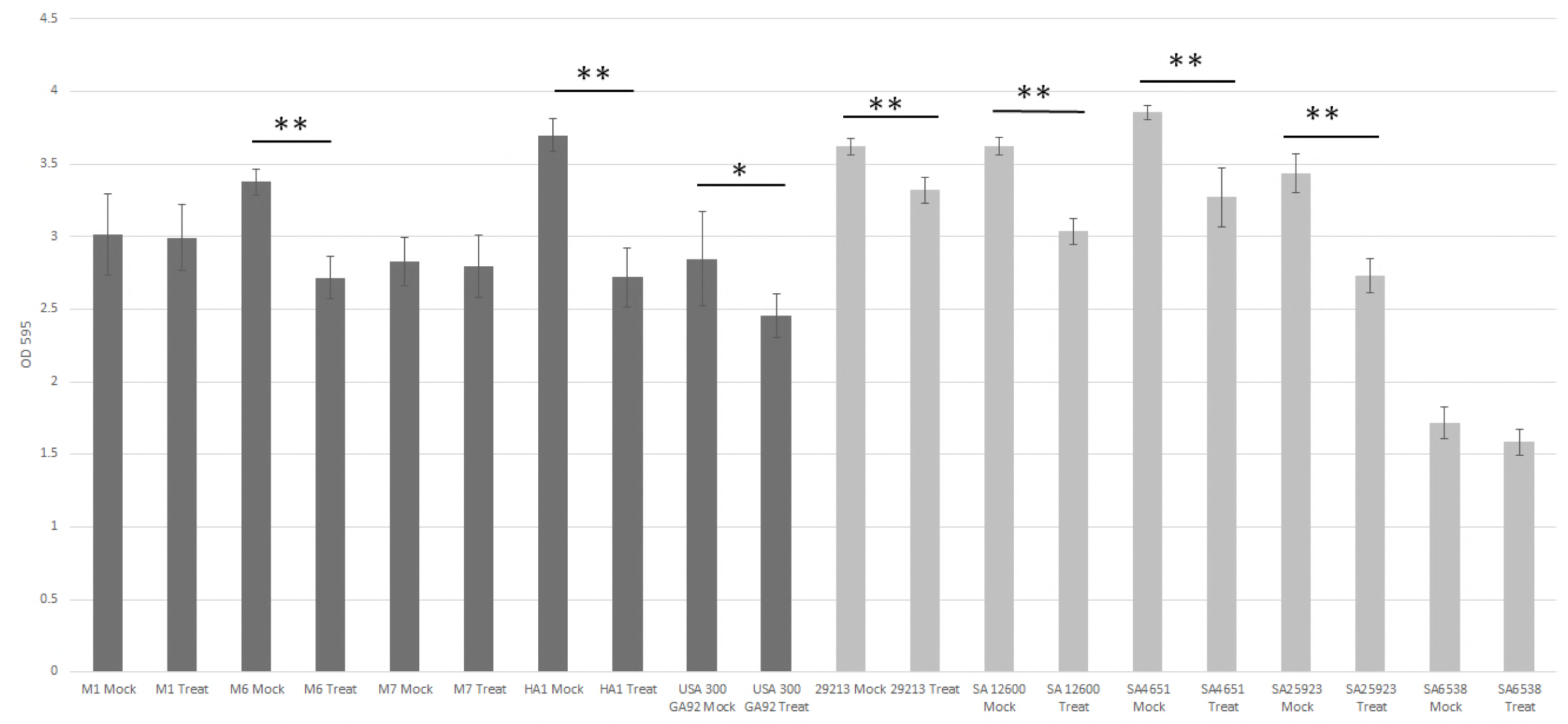
Vancomycin treatment reduces biofilm mass of certain SA strains. Established SA biofilms (24hr of growth) were treated with 2μg/mL vancomycin for 24h and then fixed and stained with crystal violet and absorbance was read at 595nm. The error bars reflect the standard error and p-values below 0.05 (*for p ≤ 0.05; ** for p ≤ 0.01) are considered significant. Dark gray are MRSA strains, light gray are MSSA strains. The controls were inoculated at the same time with a mock treatment of 210μL ddH_2_O.

### Evaluation of silver nanoparticles for anti-biofilm activity

Silver nanoparticles have anti-microbial properties due to the production of reactive oxygen species (33). Eukaryotic cells have a greater ability to neutralize reactive oxygen species as compared to prokaryotes, and so silver nanoparticles may have potential as a treatment for human infections. In order to determine the anti-biofilm effects of silver nanoparticles, biofilms were established for 24h and then 2μg/mL of 10?m silver nanoparticles was added (or citrate buffer as an untreated control; nanoparticles are suspended in this buffer). We found significant reductions in biofilm staining (Fig. 2) when analyzing all 10 SA isolates together as compared to untreated controls (mean reduction of 8.7+/-2.4%; p = 0.03). MRSA isolate biofilms showed a greater susceptibility to silver nanoparticles than MSSA strains, but the difference was not significant (p = 0.17). 3/5 MRSA isolates were susceptible (mean reduction of 11.3+/-3.4%; p = 0.01). Overall, MSSA isolate biofilms were not significantly affected by silver nanoparticle treatment; however 2/5 individual isolates showed significant reductions (mean reduction of 6.1+/-3.2%; p = 0.32).

**Figure 2:**
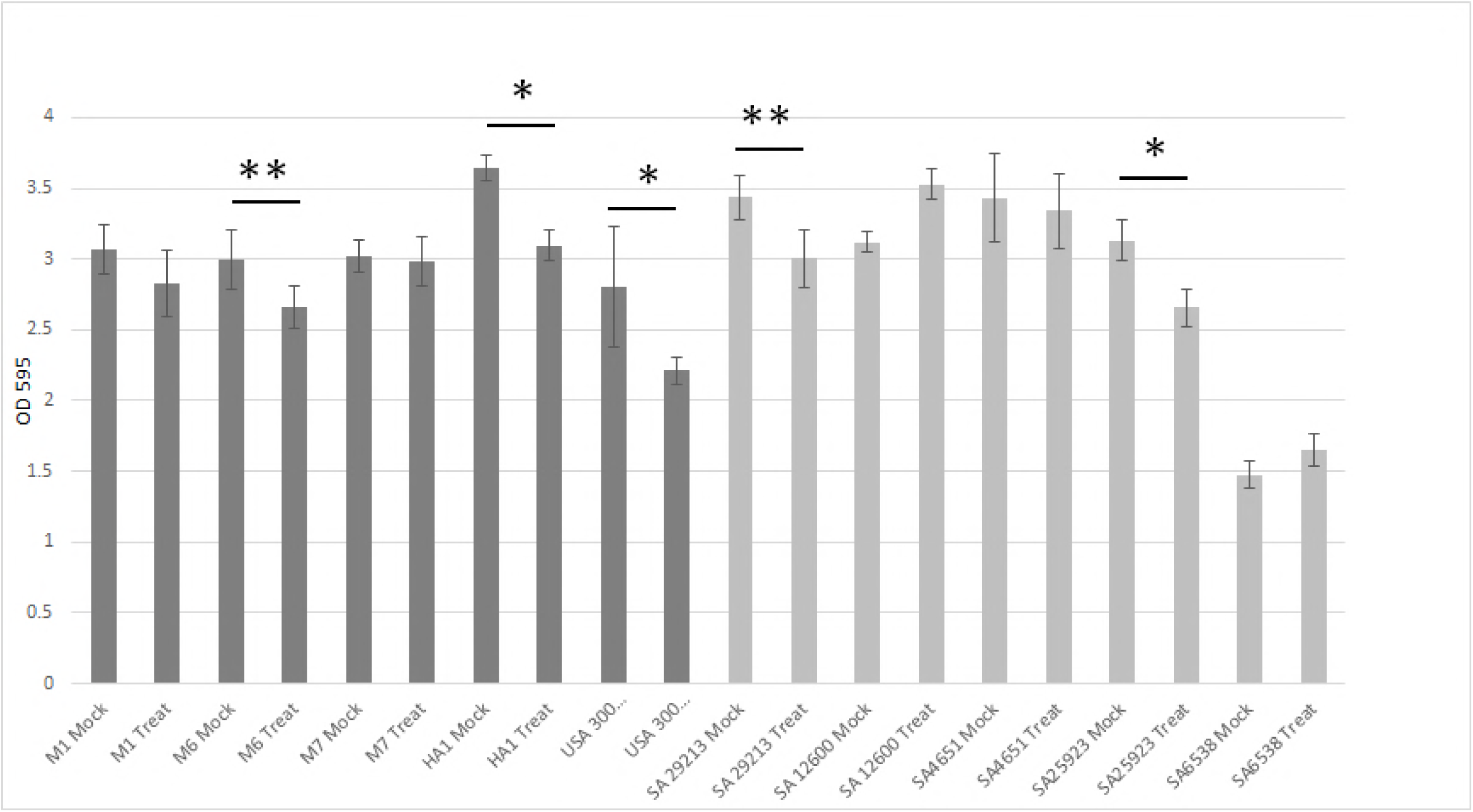
Silver nanoparticle treatment reduces biofilm mass of certain SA strains. Established SA biofilms (24hr of growth) were treated with 2μg/mL Ag nanoparticles for 24h and then fixed and stained with crystal violet and absorbance as read at 595nm. The error bars reflect the standard error and p-values below 0.05 (*for p ≤ 0.05; ** for p ≤ 0.01) are considered significant. Dark gray are MRSA strains, light gray are MSSA strains. The controls were inoculated at the same time with a mock treatment of 210μL citrate buffer.

### Synergistic effects of silver nanoparticles and vancomycin on biofilms

In order to determine if the combination of silver nanoparticles and vancomycin would have greater effects as compared to either treatment alone, we treated SA biofilms with both agents simultaneously (or citrate buffer as an untreated control; nanoparticles are suspended in this buffer) and then measured biofilm mass (Fig. 3). We found that 9/10 SA strains showed significant biofilm reductions, and with a mean reduction in biofilm mass of 20.8+/-3.8% (p = 0.0003). MRSA isolate biofilms showed a greater susceptibility to the combination treatment, with a mean reduction of 25.3+/-3.6% (p = 0.0008). MSSA strains showed a mean reduction of 16.3+/-6.4% (p = 0.04). However, there was no significant difference in MRSA vs MSSA susceptibility to the combination treatment (p = 0.17). 1 MSSA isolate (SA6538) was resistant to either treatment alone, and also resistant to the combination treatment. This isolate had the weakest baseline biofilm mass, and was somewhat susceptible to both proteinase K and DNase. We also noted that the 2 MRSA isolates that were not significantly affected by either treatment alone were susceptible to a combinatorial treatment.

**Figure 3:**
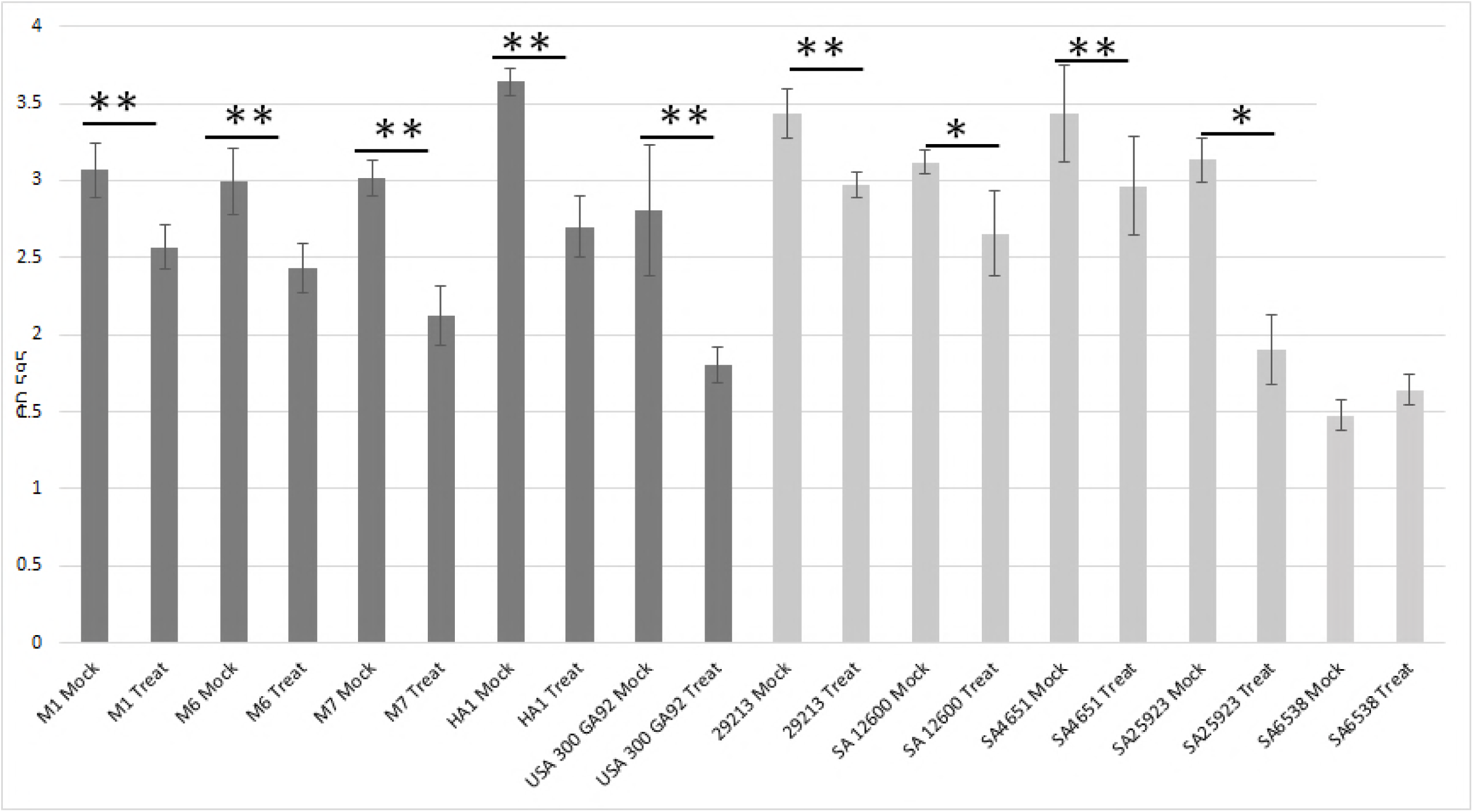
Silver nanoparticle and vancomycin combined treatment reduces biofilm of most SA strains. Established SA biofilms (24hr of growth) were treated with 2μg/mL Ag nanoparticles and 2μg/mL vancomycin for 24h and then fixed and stained with crystal violet and absorbance read at 595nm. The error bars reflect the standard error and p-values below 0.05 (*for p ≤ 0.05; ** for p ≤ 0.01) are considered significant. Dark gray are MRSA strains, light gray are MSSA strains. The controls were inoculated at the same time with a mock treatment of 210μL citrate buffer.

We then analyzed our data by strain type (MRSA, MSSA, or total SA) to determine if there were differences in susceptibility of the two many types of isolates to the three different treatments mentioned above (Fig. 4). We found that the combinatorial treatment was the most effective treatment for both MRSA and MSSA isolates, although a significant difference wasn’t always found. For MRSA isolates, the combination treatment was more effective at reducing biofilm mass than either vancomycin (p = 0.05) or silver nanoparticles alone (p = 0.01). For MSSA isolates, the combination treatment was more effective at reducing biofilm mass than silver nanoparticles alone (p = 0.04), and we also found that vancomycin treatment was more effective than silver nanoparticles alone (p = 0.05). When all SA isolates were analyzed together, we found that the combination treatment was more effective than either vancomycin (p = 0.04) or silver nanoparticles alone (p = 0.001).

**Fig 4:**
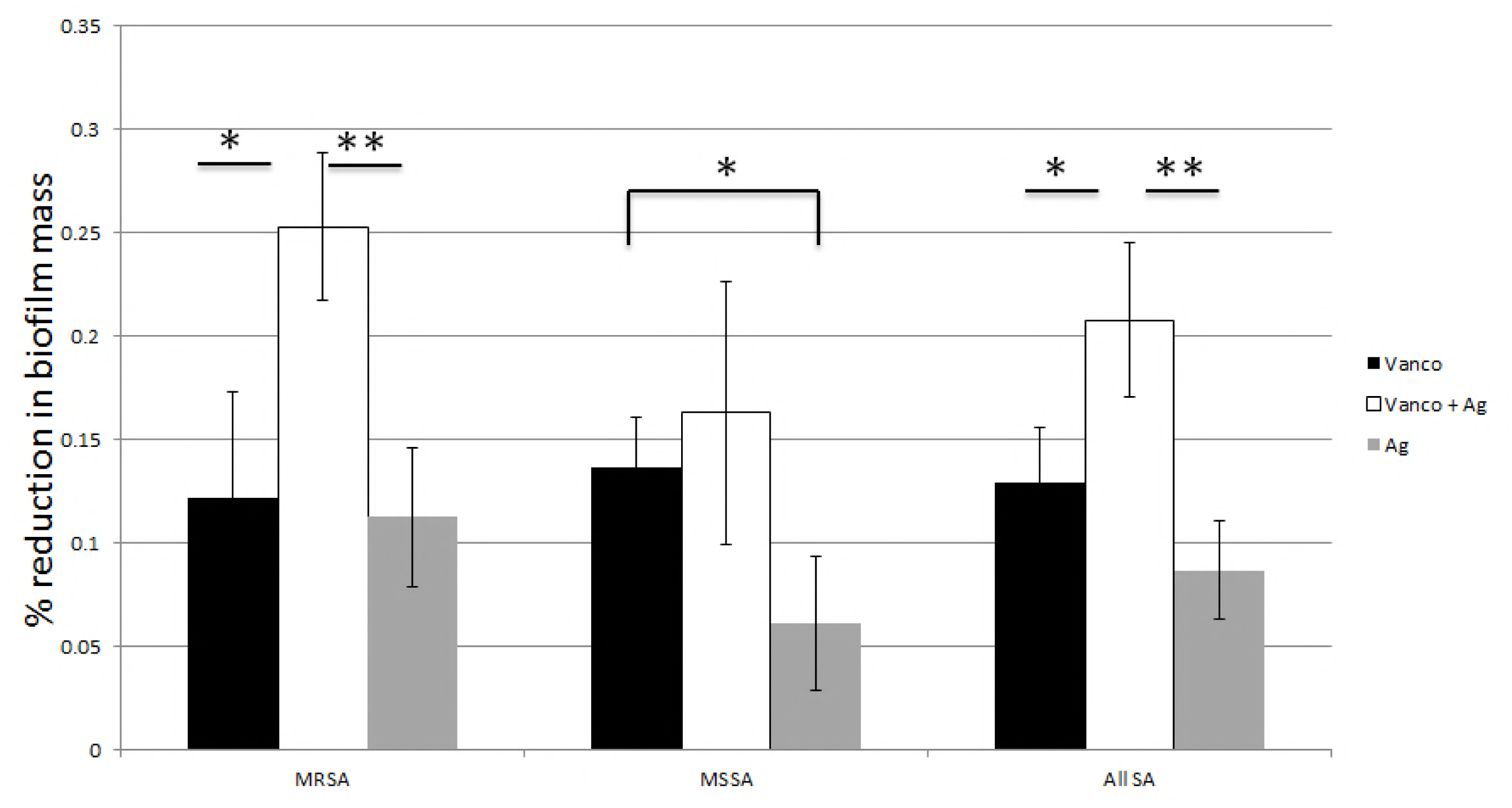
Susceptibility of SA strains to various treatments by MRSA or MSSA. The treated and control samples were compared by percent reduction after crystal violet staining and absorbance reading at 595nm. The percent reduction was measured for treatment with vancomycin alone, Ag nanoparticles and vancomycin together and Ag nanoparticles alone. The error bars reflect the standard error and p-values below 0.05 (*; ** for p ≤ 0.01) are considered significant.

## Discussion

In this study, we have characterized 10 different SA isolates for their relative biofilm-forming capacity and composition. We tested strains for their resistance to oxacillin (MRSA vs MSSA), and we also detected the presence of the *icaD* gene (involved in polysaccharide biofilm formation). We also measured the baseline biofilm mass and examined biofilm susceptibility to either proteinase K or DNase. The isolates used were quite varied in these different parameters, indicating that the isolates chosen adequately represent isolates to be encountered in clinical scenarios.

Next, we tested both an antibiotic (vancomycin) and an anti-microbial compound (silver nanoparticles) for anti-biofilm activity against a spectrum of SA isolates representing various antibiotic resistances, biofilm compositions, and relative biofilm strengths. We found that 4/5 MSSA isolate biofilms were significantly reduced following vancomycin treatment, but only 2/5 were reduced by silver nanoparticle treatment. 3/5 MRSA isolate biofilms were reduced by each treatment. 9/10 SA isolate biofilms were significantly reduced by a combinatorial treatment of vancomycin and silver nanoparticles.

Biofilms serve as defense mechanisms against immune cells and soluble immune factors, and also against drug treatment. SA biofilms tend to form canyon-like structures (10) that allow for oxygen and nutrient penetration, but are thought to prevent cells or large molecules from penetrating the biofilm. Vancomycin is a relatively large antibiotic (MW 1449), and as such is not expected to be very effective in treating SA biofilms due to poor penetration. Although 7/10 isolates tested showed significant reductions with 2μg/mL vancomycin treatment, overall the reductions were relatively minor with a mean reduction in biofilm mass of 13.0+/-2.7%. This finding may be reflective of the large size of vancomycin. We found no significant difference in vancomycin susceptibility between MRSA and MSSA strains. Susceptible isolates were *icaD*+or *icaD*-, had with high or low scores in the congo red agar test, showed either strong or weak biofilm staining characteristics in a crystal violet assay, and their biofilms were susceptible to proteinase K or DNase treatments. These results suggest that vancomycin has broad spectrum activity against SA biofilms, but overall the reduction in biofilm mass is relatively minor in a tissue culture test.

Silver nanoparticles are known to have toxic properties when exposed to cells. The proposed mechanism is via the formation of reactive oxygen species as silver ions are released. Silver nanoparticles have been explored for activity against a number of pathogenic bacteria, including SA (34, 35). They have also been shown to be able to prevent SA biofilm formation (36, 37). However, their activity against established SA biofilms has not been extensively explored. One report showed that they are effective against *Pseudomonas aeruginosa* biofilms (38). In that study, they examined biofilm structures following silver nanoparticle treatment and found that 10nm nanoparticles penetrate into the biofilm matrix more effectively than other sizes, resulting in greater activity of aztreonam against lipid bilayers in the target cells.

We found that silver nanoparticle treatment of established SA biofilms has activity against both MRSA and MSSA isolates, but that more isolates are unaffected by this treatment (5/10 affected) as compared to vancomycin (7/10 affected). No clear patterns emerged amongst the isolates that were resistant to silver nanoparticle treatment; these isolates could have a strong or a weak base biofilm mass, they could be resistant to proteinase K treatment or DNase treatment, and some strains were equally susceptible to proteinase K or DNase treatments.

Our final set of tests was a combinatorial treatment of silver nanoparticles and vancomycin. We found that 9/10 SA isolates showed significant reductions in biofilm staining following this dual treatment, suggesting that the two treatments work in concert to eliminate biofilm mass. The dual treatment was more effective than either treatment alone for the group of MRSA isolates, and also for all SA isolates, but not for the group of MSSA isolates alone.

Interestingly, we noted that the same two MRSA isolates (M1 and M7) that showed biofilm resistance to vancomycin treatment were also resistant to silver nanoparticle treatment. These two isolates had similar effects following treatment with proteinase K or with DNase, and also had similar baseline biofilm staining results (see Table 1). Strain M7 lacked the *icaD* gene, while strain M1 contained this gene, and both strains gave similar colony morphologies on congo red agar (+++). 1 MSSA isolate (SA6538) was resistant to both singular treatments, and was also the only strain resistant to the combinatorial treatment. This isolate had a weak baseline biofilm mass, was++ on the congo red agar test, lacked the *icaD* gene, and was somewhat susceptible to both proteinase K and DNase. Interestingly, although this strain had the weakest biofilm mass of any isolate tested, it did have similar susceptibility to proteinase K and DNase treatments when compared to strains M1 and M7.

SA biofilms have clinical relevance in terms of both transmission, and *in vivo* pathogenesis after host invasion. These findings could be useful both to prevent transmission via treatment of catheter materials, and also for treatment of established biofilms *in vivo*. Future directions could include analysis of other antibiotics, such as those with a smaller molecular weight. Other types of metal nanoparticles, or nanoparticles of varying sizes, could also be explored for anti-SA biofilm activity.

## Materials and Methods

### Isolation and sources of SA

Nine SA strains were used in this study. Strains M1, M6 and M7 were isolated from a sports training facility in Utah. Strain HA1 is a local hospital-associated isolate. Strain USA300-GA92 was acquired from BEI resources (Manassas, VA,), and strains SA29213, SA12600, SA4651, and SA25923 were purchased from American Type Culture Collection (Manassas, VA).

### Characterization of biofilms by congo red agar colony morphology

37g Brain Heart Infusion, 0.8g congo red, 36g sucrose, and 12g agar were mixed in 1L and autoclaved (39). SA strains were streaked to single colonies and grown at 37°C for 48h. The resulting growth was then categorized according to color as-,+,++, or+++; with-being bright red colonies with no black around the colonies;+being mostly red with black colonies only in high-density areas,++ being almost entirely black with some red colonies, and+++being all colonies being black (40).

### *icaD* genotyping

DNA was extracted from SA strains (41), and the *icaD* gene was detected via PCR. Primers used were F: (5′-GAACCGCTTGCCATGTGTTG-3′) and R: (5′-GCTTGACCATGTTGCGTAACC-3′) giving a product of 483 bp as described by (42), and confirmed with the control strain USA300 0114. Thermal cycling parameters were followed as described by each source with the exception that we performed 40 cycles of the amplification step in each reaction. We used a standardized reaction mixture of 25μl, consisting of 15.8μl ddH_2_O, 2.5μl of PCR buffer (15mM MgCl_2_), 2.5μl dNTPs (2.5mM), 1μl of each primer (5μM stock), and 0.2μl of AmpliTaq DNA polymerase from (5U/μl) supplied by Applied Biosystems. PCR products were visualized under UV light on a 1% agarose gel stained with ethidium bromide.

### Crystal Violet biofilm assay

Biofilm mass was measured using an adaptation of a previously described assay using 96-well plates (43, 44). Strains to be tested were grown overnight at 37° C in tryptic soy broth (TSB) (Sigma-Aldrich, St. Louis, MO). Cultures were then diluted 1:200 in 66% TSB with 0.5% glucose added. 200μL of culture dilution were added to the wells of a 96-well, flat-bottomed, tissue-culture plate (Falcon #353916, Corning Incorporated, Corning, NY) in quadruplicate, covered, and incubated for 24 hours at 37° C. The liquid and non-adhered cells were then removed from the wells by gently overturning the plate onto paper towels. Each well was then gently washed with 1x phosphate-buffered saline (PBS) and allowed to dry. Once dry, 205μL of 100% ethanol was added to each well to fix the biofilms, incubated for 15 minutes at room temperature, and then emptied onto a paper towel. Once the ethanol had dried 205μL of 0.1% crystal violet dye was added to each well and incubated for 15 minutes at room temperature. The dye was then emptied as before, by overturning the plate onto paper towels, and three washes with 210μL of ddH_2_O were performed, dumping the water between each wash. Once dry, stain was eluted from the biofilms with 205μL of a mixture of 1/3 volume of EtOH with 40 mM HCl and 2/3 volume of acetone added to each well. The wells were sealed and incubated with this solution for 15 minutes at 37° C with 100 rpm shaking. 80μL of eluted stain was removed from each well and transferred to a new plate for reading and the absorbance at 595nm was measured for each well. All crystal violet biofilm measurements in these studies were conducted in quadruplicate, and then repeated again in quadruplicate, for a total of 8 independent measurements.

### Treatments to disrupt biofilms

The biofilms used for this test were grown as previously described in the crystal violet biofilm assays. After a period of 24h growth at 37°C the liquid contents of the wells were emptied onto paper towels. This was done in one quick, yet gentle motion to preserve the biofilm structure. When the wells were fully emptied each well designated for one of three treatments was inoculated with 210μL of either vancomycin at a concentration of 2μg/mL or 10nm silver nanoparticles at a concentration of 2μg/mL in sodium citrate or a combination treatment of 2μg/mL vancomycin and 10nm silver nanoparticles at 2μg/mL. The controls were also inoculated at this same time with a mock treatment of 210μL ddH_2_O to control for the vancomycin treatment and 210μL of sodium citrate at a concentration of 2mM to control for the silver nanoparticle treatment and also for the combination treatment. The samples were incubated at 37°C for 24 hours. At the end of the 24h period the wells were again emptied by dumping the contents onto paper towels and were left upside down for approximately 24-48h, or until completely dried out. Once dry, the biofilms were fixed, stained, washed and eluted as mentioned in the Crystal Violet biofilm assay section.

### Statistical analysis

Paired, one-tailed student’s t tests were employed to determine statistical significance.

## Acknowledgements

This research was supported by a BYU Turkey Vaccine Award to BKB, and a BYU ORCA award to BBH. The funders had no role in study design, data collection and interpretation, or the decision to submit the work for publication.

We are grateful to Richard Robison (Brigham Young University) and Russell Osguthorpe (Primary Children’s Hospital, Salt Lake City, UT) for sharing human MRSA isolates. Some SA isolates were obtained from the American Type Culture Collection and from BEI Resources.

